## Supplementary Figures for "An integrated resource for functional and structural connectivity of the marmoset brain"

[1] Department of Neurobiology, University of Pittsburgh Brain Institute, University of Pittsburgh, Pittsburgh PA 15261, USA
[2] Center for Excellence in Brain Science and Intelligence Technology, Institute of Neuroscience, CAS Key Laboratory of Primate Neurobiology, Chinese Academy of Sciences, Shanghai, China
[3] Laboratory of Neuroinformatics, Nencki Institute of Experimental Biology of the Polish Academy of Sciences, 02-093 Warsaw, Poland
[4] Department of Physiology and Neuroscience Program, Biomedicine Discovery Institute, Monash University, Clayton, VIC 3800, Australia
[5] Center for Brain and Cognition, Computational Neuroscience Group, Department of Information and Communication Technologies, Universitat Pompeu Fabra, Roc Boronat 138, Barcelona, 08018, Spain
[6] Universidad de San Andrés, Vito Dumas 284 (B1644BID), Buenos Aires, Argentina
[7] Cerebral Microcirculation Section, Laboratory of Functional and Molecular Imaging, National Institute of Neurological Disorders and Stroke, National Institutes of Health (NINDS/NIH), Bethesda, MD 20892, USA
[8] Department of Neurobiology, Affiliated Mental Health Center & Hangzhou Seventh People's Hospital, Zhejiang University School of Medicine
[9] MOE Frontier Science Center for Brain Science and Brain-machine Integration, Zhejiang University, Hangzhou, China.
[10] Scientific and Statistical Computing Core, National Institute of Mental Health, National Institutes of Health (NIMH/NIH), Bethesda, MD 20892, USA
[11] Institució Catalana de la Recerca i Estudis Avançats (ICREA), Passeig Lluís Companys 23, Barcelona, 08010, Spain
[12] Department of Neuropsychology, Max Planck Institute for Human Cognitive and Brain Sciences, 04103 Leipzig, Germany
[13] School of Psychological Sciences, Monash University, Melbourne, Clayton VIC 3800, Australia
[14] Shanghai Center for Brain Science and Brain-Inspired Intelligence Technology, Shanghai, China
[15] Lingang Laboratory, Shanghai 200031, China
[16] University of Chinese Academy of Sciences, Beijing, China
* Corresponding authors

### Supplementary Figures

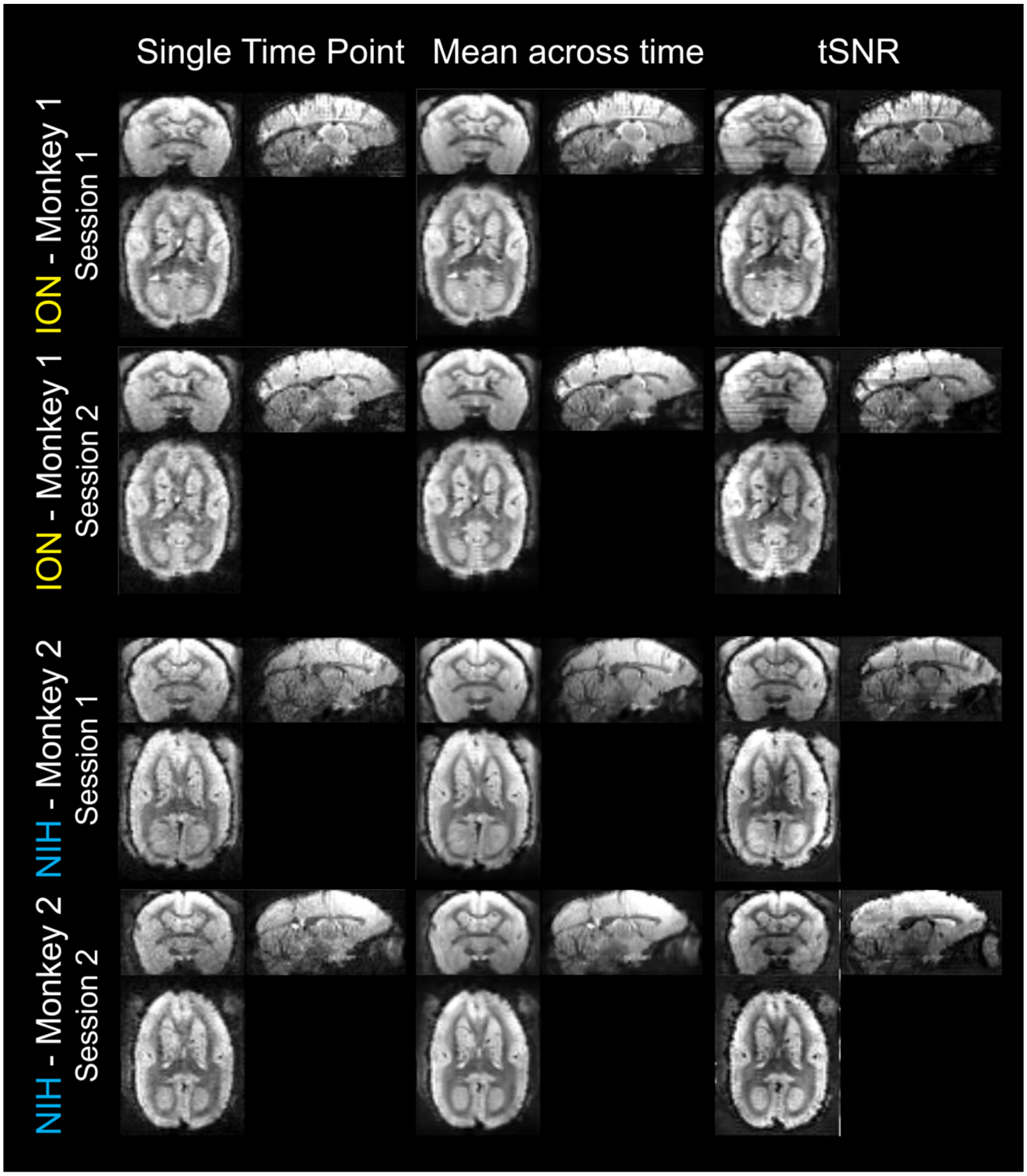


**Supplementary Figure 1. Example images of the ION and the NIH datasets.** Single time points, mean images (averaged across time for one fMRI run), and tSNR images (calculated from one fMRI run) are presented for four sessions of two flagship monkeys from ION and NIH, respectively. The tSNR image of each session was calculated by 3dTstat of AFNI.

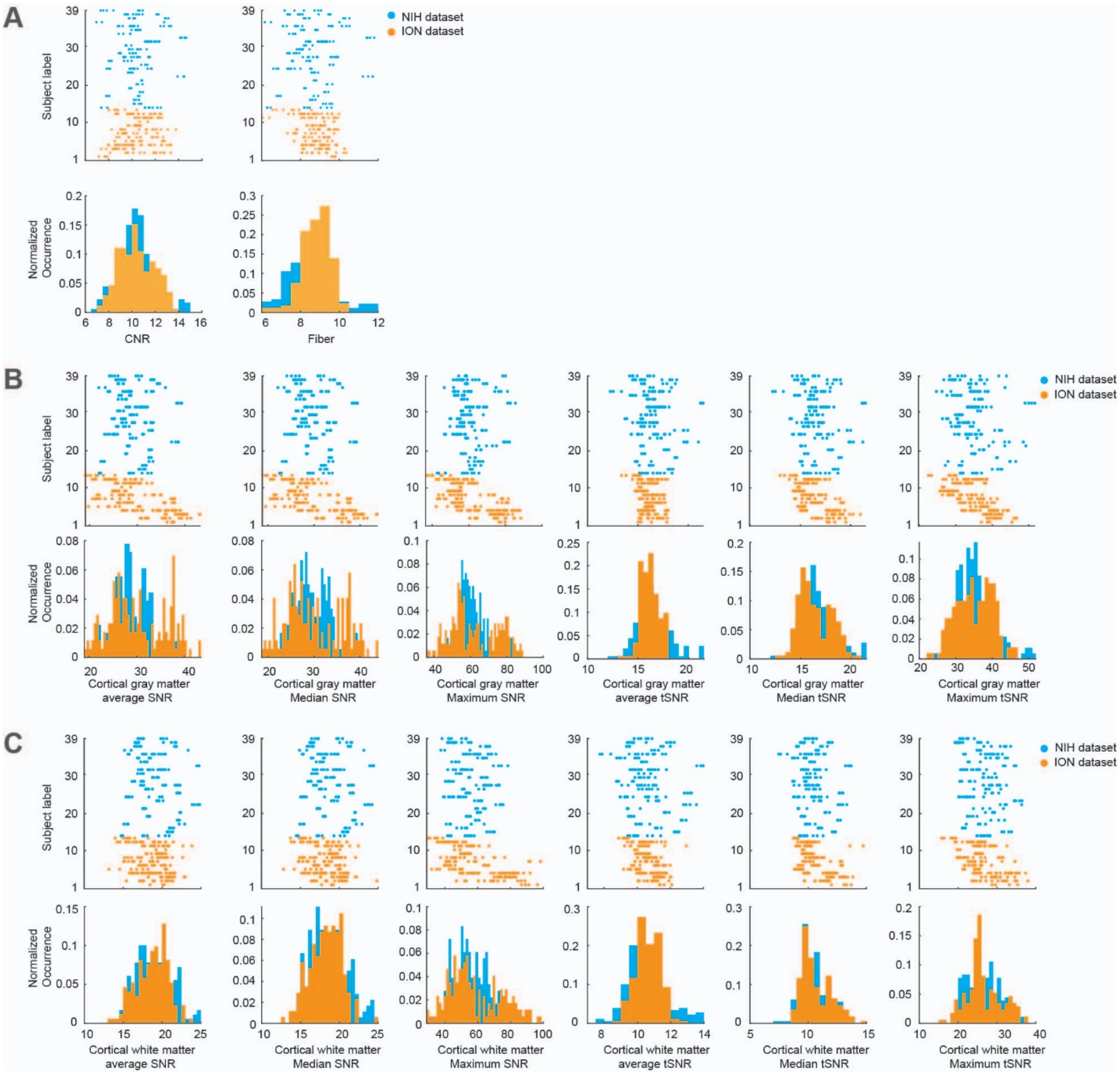


**Supplementary Figure 2. Similar quality measurements of the ION and the NIH datasets.** (A) The raster plots and their histograms present the CNR (Contrast to Noise Ratio: the mean of the gray matter intensity values minus the mean of the white matter intensity values divided by the standard deviation of the values outside the brain) and the Fiber (Foreground to Background Energy Ratio: the variance of voxels inside the brain divided by the variance of voxels outside the brain) of two datasets (the blue represents the results from the NIH dataset, and the yellow represents the results from the ION dataset); the results of the two-sided Wilcoxon rank test between two datasets (N-NIH =180 N-ION =172) are p=0.45 and p=0.11, respectively. (B) the raster plots and histograms present the average SNR, median SNR and max SNR, average tSNR, median tSNR, and max tSNR of the cortical gray matter from two datasets (N-NIH =180 N-ION =172). The two-sided Wilcoxon rank tests for SNR are p=0.259, p=0.824, and p=0.968, and for tSNR are p=0.435, p=0.625, and p=0.2, respectively. (C) presents the average SNR, median SNR and max SNR, average tSNR, median tSNR, and max tSNR of cortical white matter from two datasets (N-NIH =180 N-ION =172). The two-sided Wilcoxon rank tests for SNR are p=0.712, p=0.32 and p=0.42, and for tSNR are p=0.062, p=0.086, and p=0.908, respectively. The NIH and the ION datasets have no significant difference in the above quality assurance (QA) measurements. The tSNR image of each session was calculated by 3dTstat of AFNI. The SNR and tSNR values were calculated by "the mean value of gray matter voxels divided by the standard deviation of background noises."

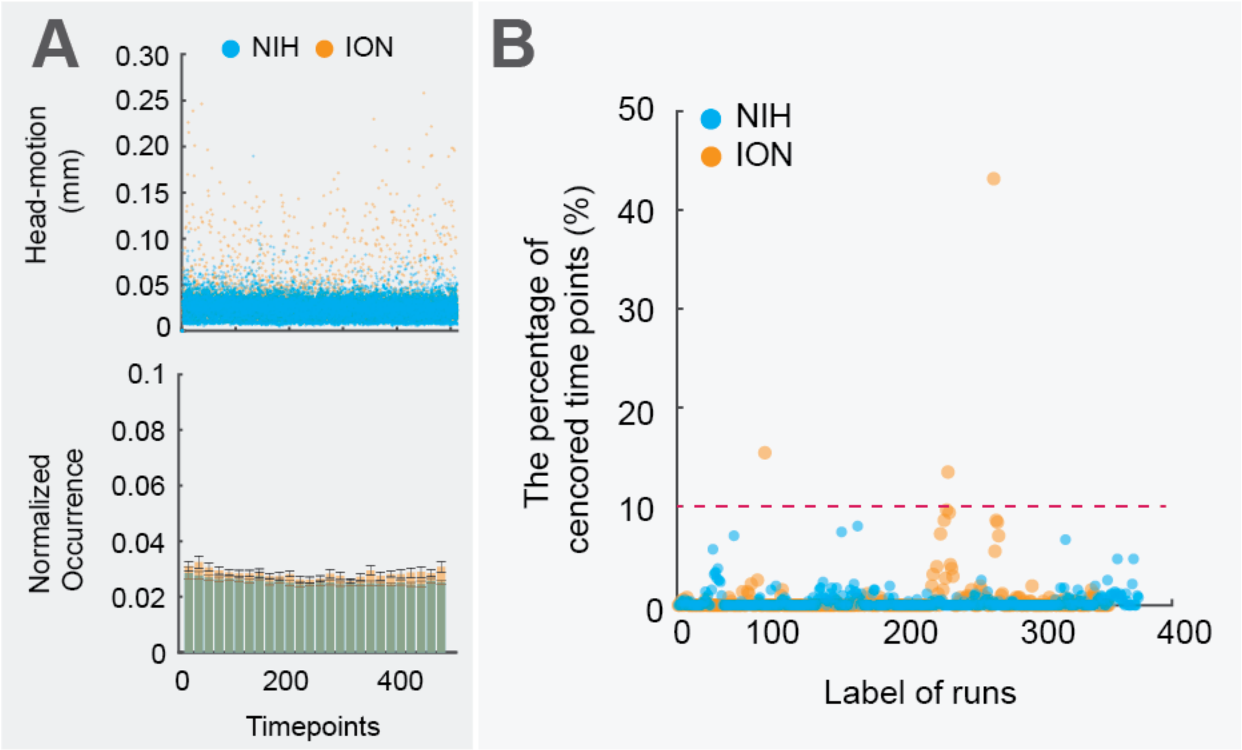


**Supplementary Figure 3. Head motions of the ION and the NIH datasets.** (A) the top panel presents head-motion (weighted euclidean norm of six motion parameters) across 512 timepoints of different datasets (the blue represents the NIH dataset, N_subject=26, and the yellow is the ION dataset, N_subject=13). Each dot is the head-motion measure of each fMRI at a one-time point. The bottom panel presents the corresponding histogram statistics from each dataset (bin size = 20 timepoints, the bar charts are presented as mean values +/- 95% C.I., average normalized occurrence value across 512 timepoints is 0.03), which indicates that head-motion levels are similar across datasets. (B) The percentage of censored time points (motion > 0.2mm and temporal outlier > 0.1) for each fMRI. Most animals and fMRI runs (710 runs) have low head motion and censored time points, suggesting the effectiveness of our head-constrained and training approaches. Note that the three fMRI runs with extensive head motions (more than 10% time points were censored) were excluded from our analysis. However, we still included those three runs in the source (raw) data release.

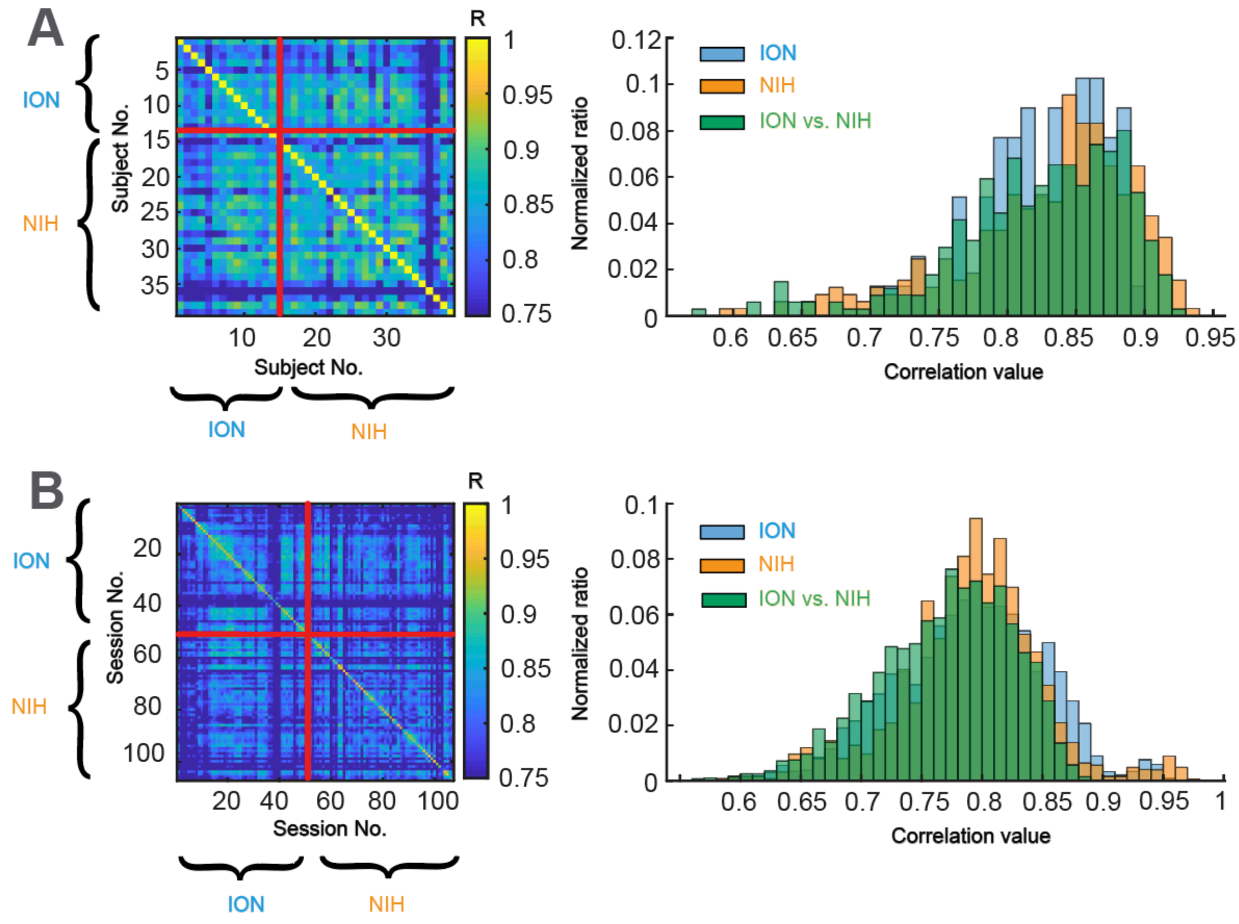


**Supplementary Figure 4. Similar measurements of the whole-brain functional connectivity across subjects and sessions from ION and the NIH datasets.** (A) The left heatmap represents the functional connectivity similarity matrix across subjects (we used the 2D Pearson correlation coefficient as the similarity metric). Subjects No. 1-13 come from the ION dataset, and the rest subjects (No. 14-39) are from the NIH dataset. To demonstrate the high similarity across sites, we further made a histogram plot of the similarity metric for the subjects from the ION dataset (light blue), the NIH dataset (orange), cross ION-NIH dataset (Green). They present high similarity (One-way ANOVA multiple comparison tests, df=2, F=0.92, p = 0.4008). (B) Same as (A), we present the functional connectivity similarity matrix across sessions, Sessions No. 1-55 come from the ION dataset, and the rest of the sessions (No. 56-107) are from the NIH dataset. They present high similarity (One-way ANOVA multiple comparison tests, df=2, F=97.63, p = 1.717)

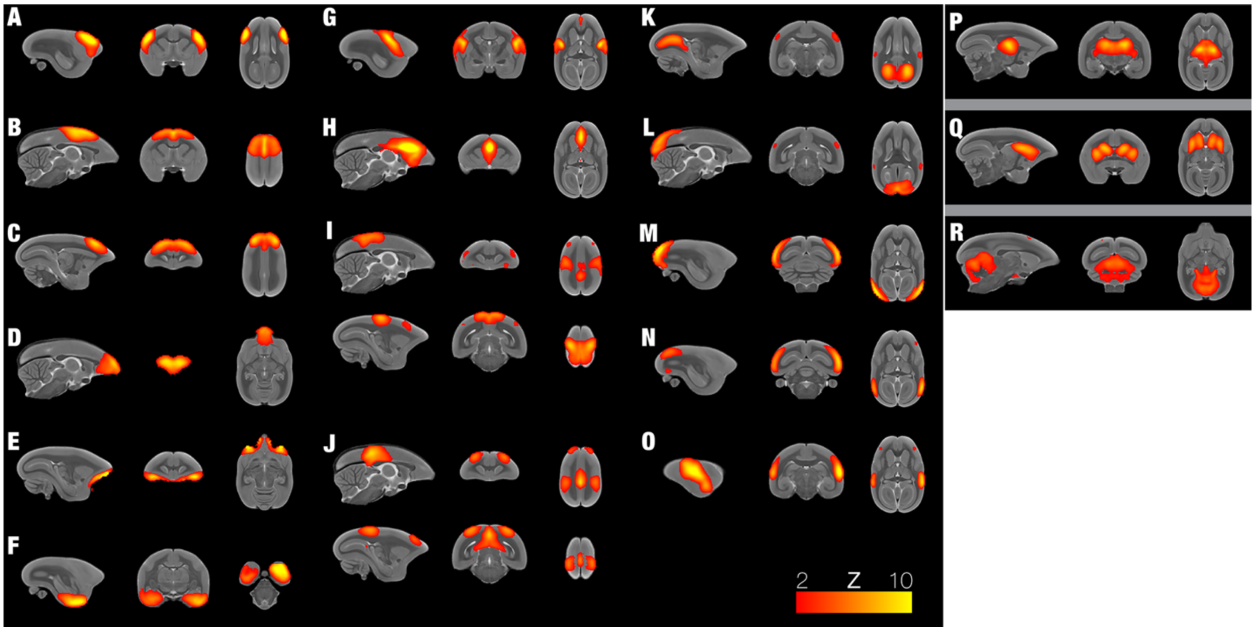


**Supplementary Figure 5. Identified resting-state functional networks.** We found 18 networks by group-ICA analysis (A-O cortical networks; P-R subcortical networks), including (A) the ventral somatomotor, (B) the dorsal somatomotor, (C) the premotor, (D) the frontal pole, (E) the orbital frontal cortex, (F) the parahippocampus/temporal pole, (G-H) the salience-related network, (I-J) two trans-modal networks, which are frontoparietal (I) and the default-mode-network-related (J), the visual-related networks from primary visual cortex (K-M) to functional higher-level regions (N-O) and subcortical networks, the thalamus (P), the striatum (Q), and the cerebellum (R).

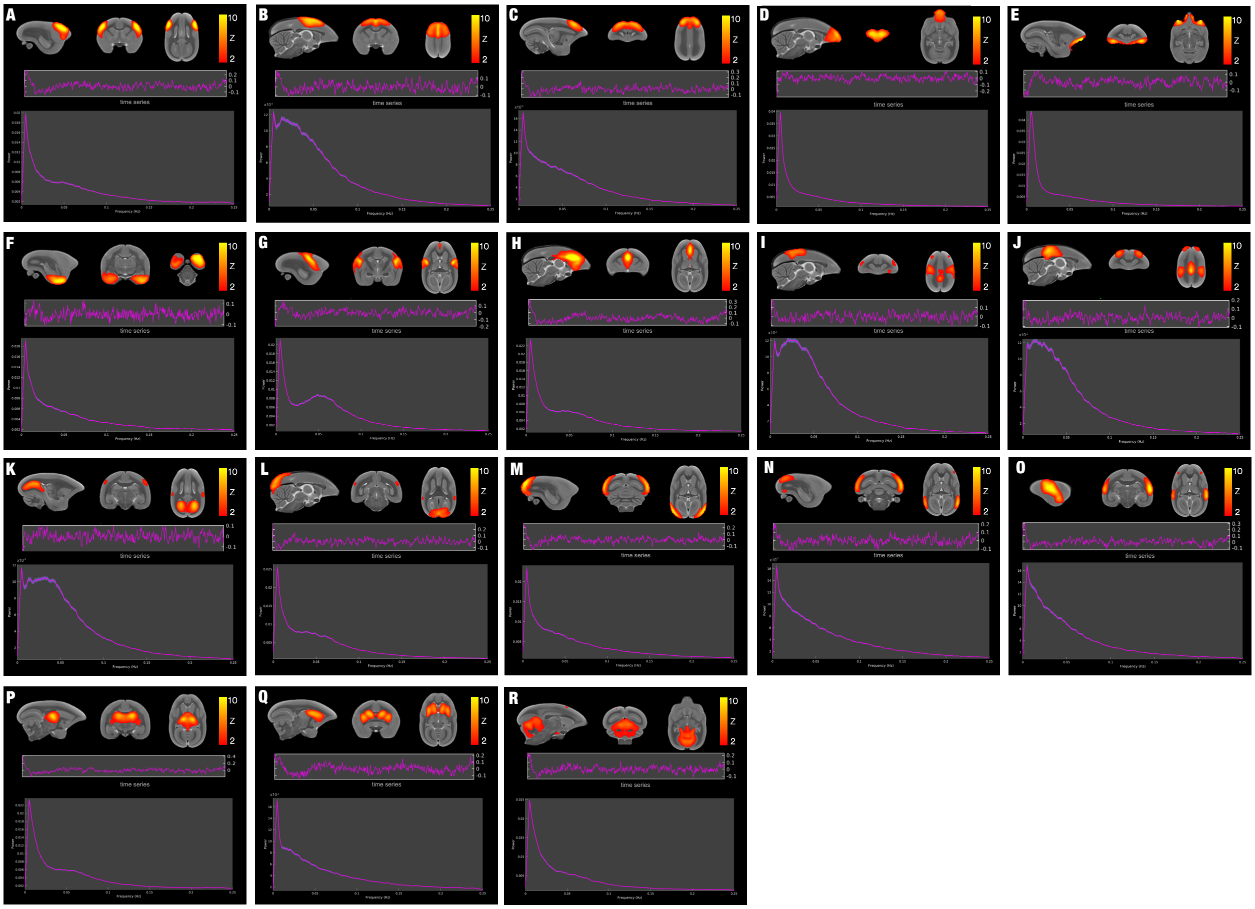

**Supplementary Figure 6. Similar to Figure S5 but with the time series and frequency power plotted for each component.** The networks include: (A) the ventral somatomotor, (B) the dorsal somatomotor, (C) the premotor, (D) the frontal pole, (E) the orbital frontal cortex, (F) the parahippocampus/temporal pole, (G-H) the salience-related network, (I-J) two trans-modal networks, which are frontoparietal (I) and the default-mode-network-related (J), the visual-related networks from primary visual cortex (K-M) to functional higher-level regions (N-O) and subcortical networks, the thalamus (P), the striatum (Q), and the cerebellum (R).

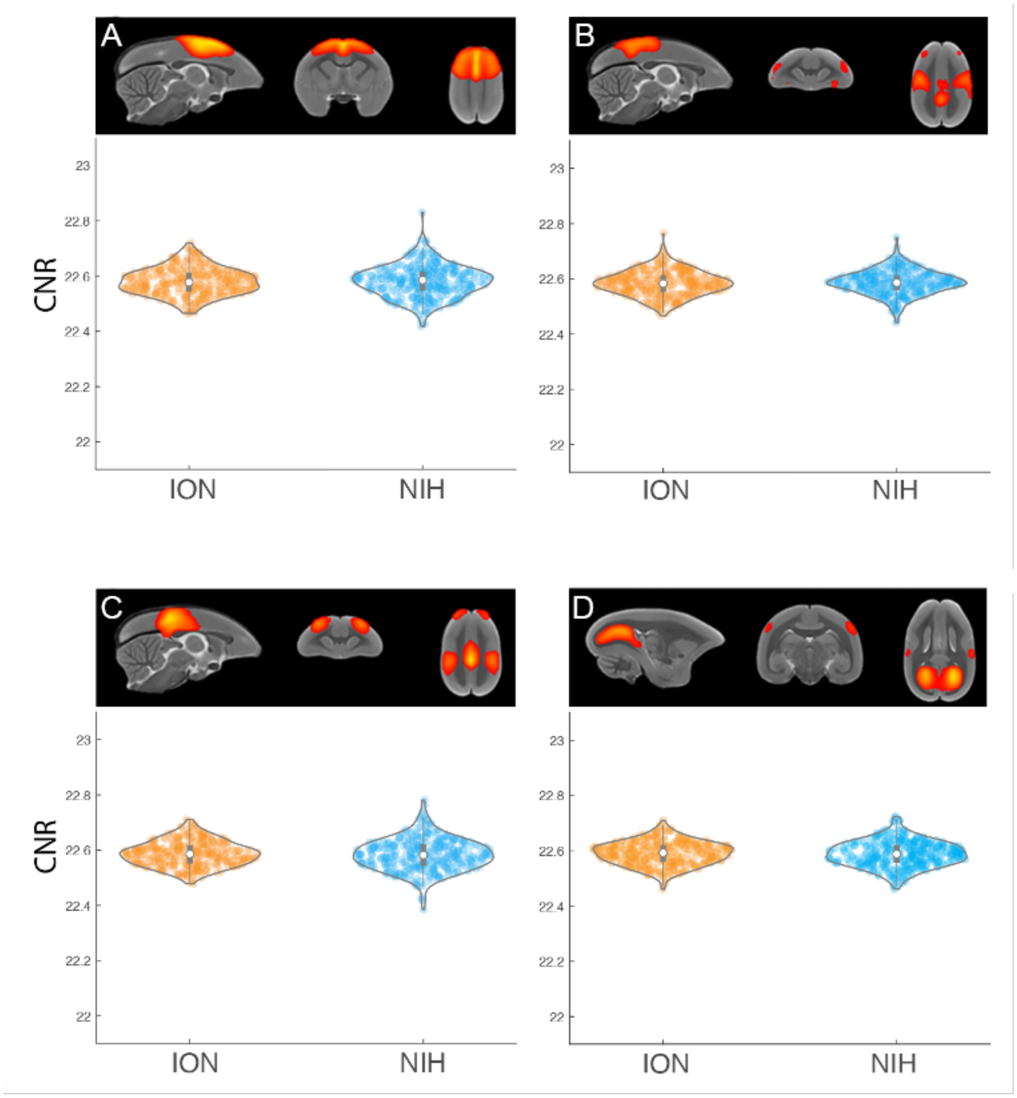


**Supplementary Figure 7. Similar temporal CNR (tCNR) of the ION and the NIH datasets.** Four networks (A) the dorsal somatomotor, (B) frontoparietal network, (C) default-mode-network, and (D) primary visual network are shown. The tCNR is the variance of optimal resting-state fMRI components after ICA contrast to the noise components). We selected four ICA components from network results, Data are presented by the violin and the box plots in which the white point represents the average value (B: the dorsal somatomotor, ION dataset N=323 sessions: 25th percentile 22.565 and 75 percentile 22.62, average value 22.594, maximum: 22.71; minimum 22.468; NIH dataset N=368 sessions: 25th percentile 22.554 and 75 percentile 22.613, average value 22.584, maximum: 22.764; minimum 22.455; I: the frontoparietal, ION dataset N=323 sessions: 25th percentile 22.56 and 75 percentile 22.62, average value 22.593, maximum: 22.71; minimum 22.46; NIH dataset N=368 sessions: 25th percentile 22.557 and 75 percentile 22.62, average value 22.585, maximum: 22.727; minimum 22.451; J: the default-model frontoparietal, ION dataset N=323 sessions: 25th percentile 22.562 and 75 percentile 22.62, average value 22.588, maximum: 22.716; minimum 22.463; NIH dataset N=368 sessions: 25th percentile 22.515 and 75 percentile 22.614, average value 22.568, maximum: 22.827; minimum 22.367. K: the visual-related network, ION dataset N=323 sessions: 25th percentile 22.558 and 75 percentile 22.62, average value 22.589, maximum: 22.749; minimum 22.425; NIH dataset N=368 sessions: 25th percentile 22.534 and 75 percentile 22.62, average value 22.574, maximum: 22.85; minimum 22.316) and compared their temporal CNR between the two datasets. No significant difference was found between the two sites/scanners (The two-sided Wilcoxon rank tests, $p>0.05$, N-ION=346, N-NIH=364).

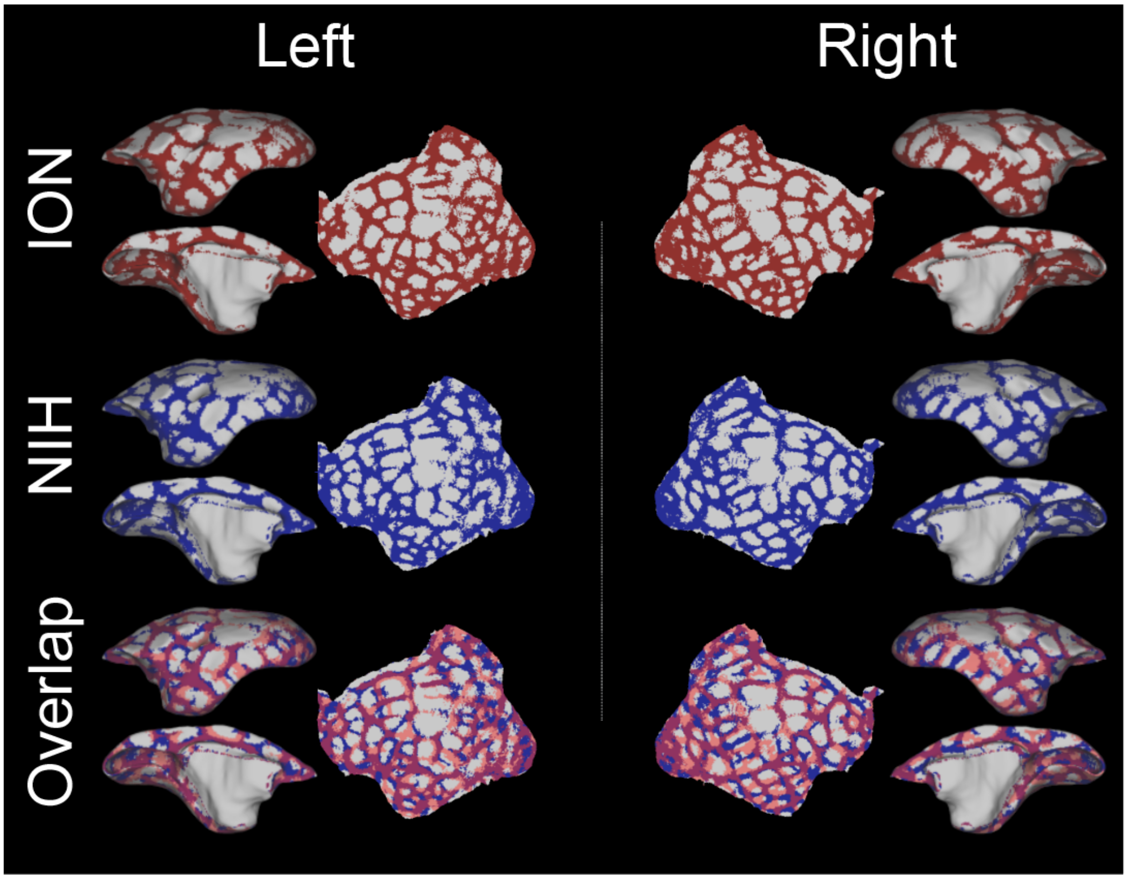


**Supplementary Figure 8. Boundary maps in both hemispheres from NIH and ION Dataset are highly similar.** Top and middle panel: The boundary maps in both hemispheres from NIH and ION datasets after thresholding both at the 75th percentile of boundary map values. Bottom: The comparison between two boundary maps (Top and middle) in both hemispheres. Light blue: NIH boundaries; pink: ION boundaries; purple: the overlapping boundaries between datasets.

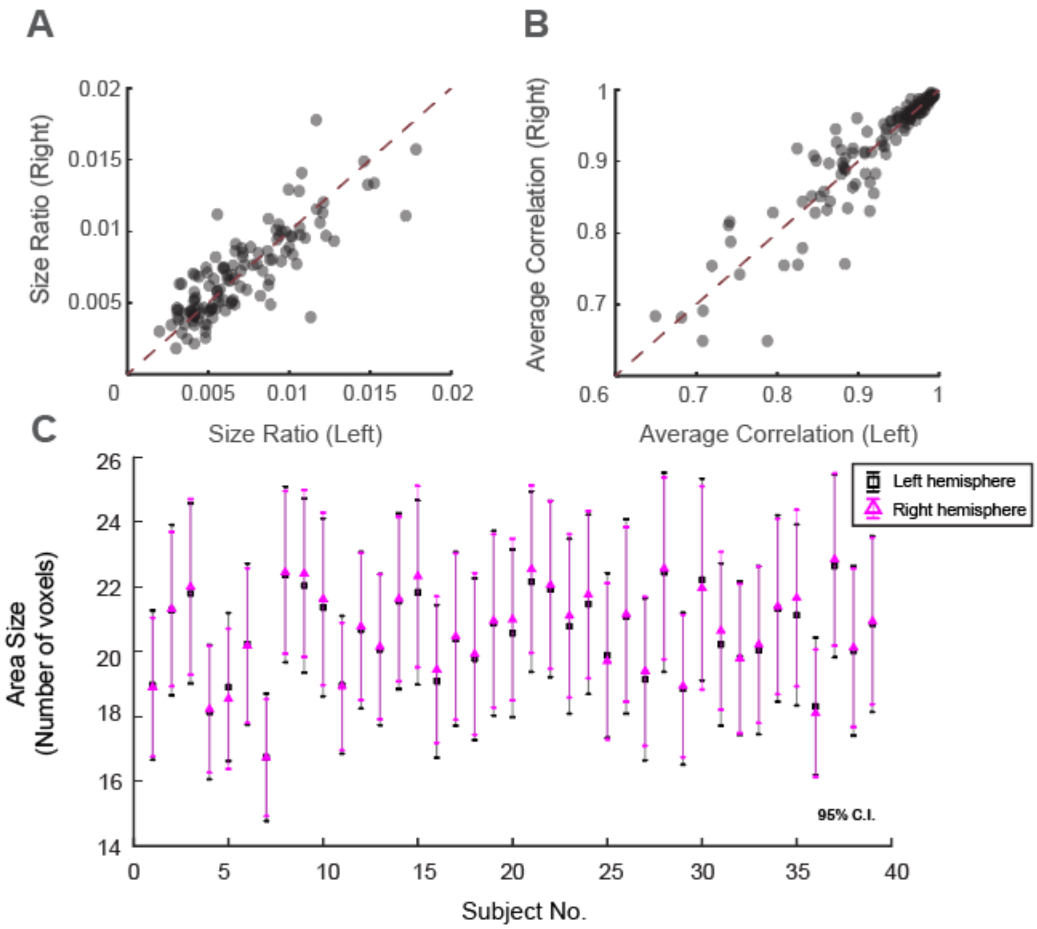


**Supplementary Figure 9. The functional parcels are highly similar across the hemisphere.** (A) All parcel sizes. (B) The functional connectivity patterns of all parcels. The dashed line represents the diagonal line. (C) The surface area size in each hemisphere by inverted warping of the standard average surfaces to every subject's native space (the number of areas in each hemisphere is 96), representing the data by the mean +/- 95% confidence interval.

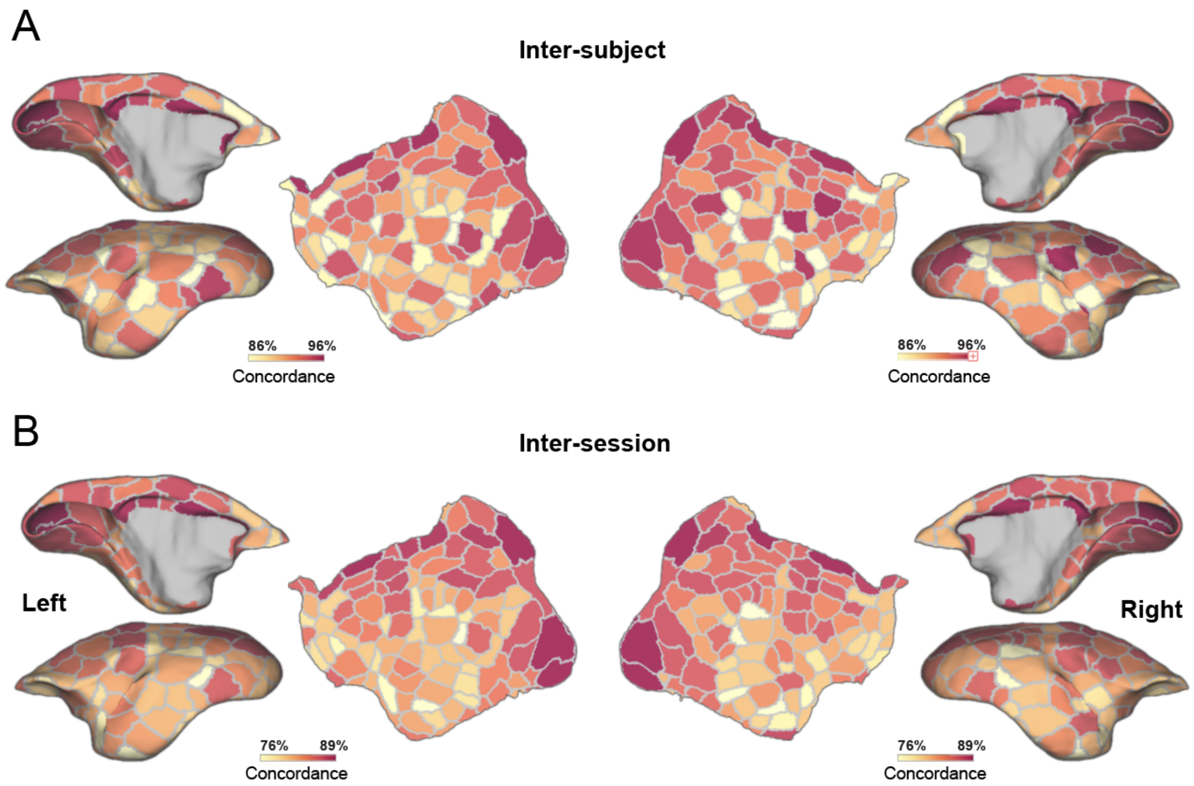


**Supplementary Figure 10. The variation of individual mapping parcels by the deep neural network.** (A) The concordance of inter-subject parcels. (B) The concordance of inter-session parcels.

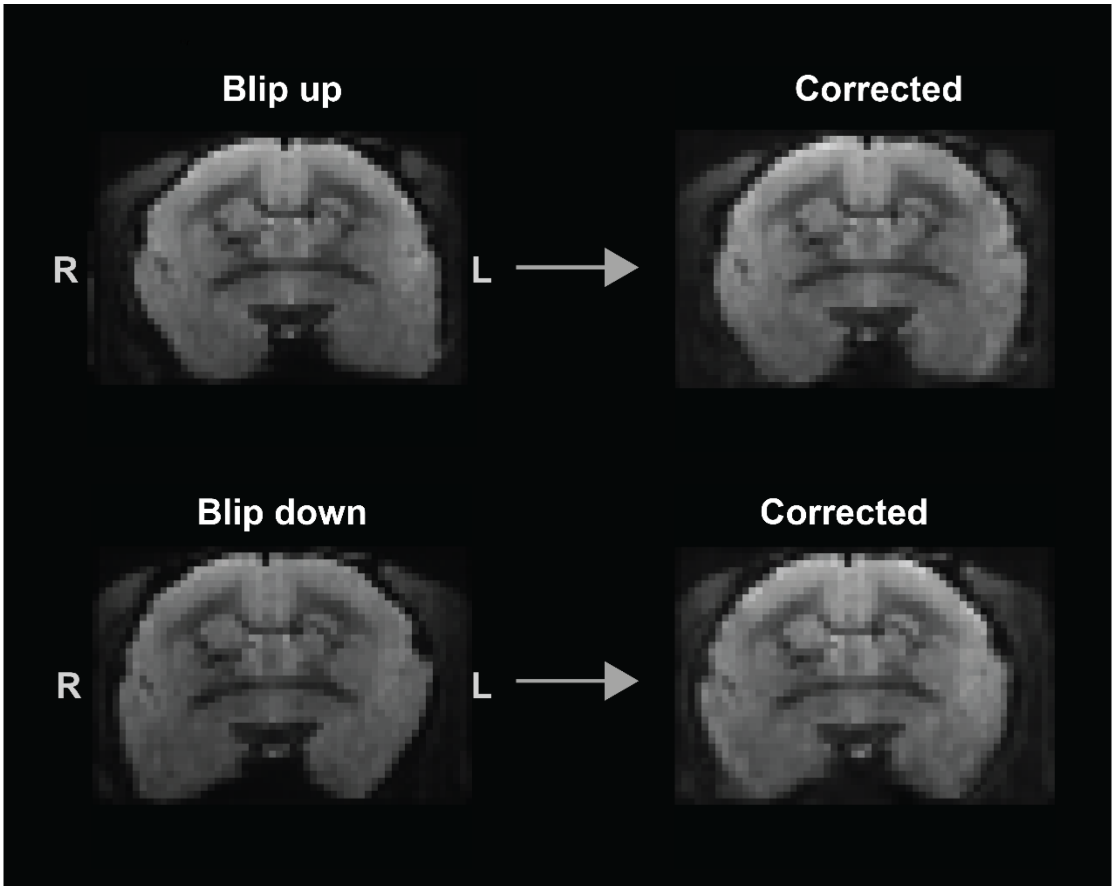


**Supplementary Figure 11. Examples of the top-up EPI distortion correction.**

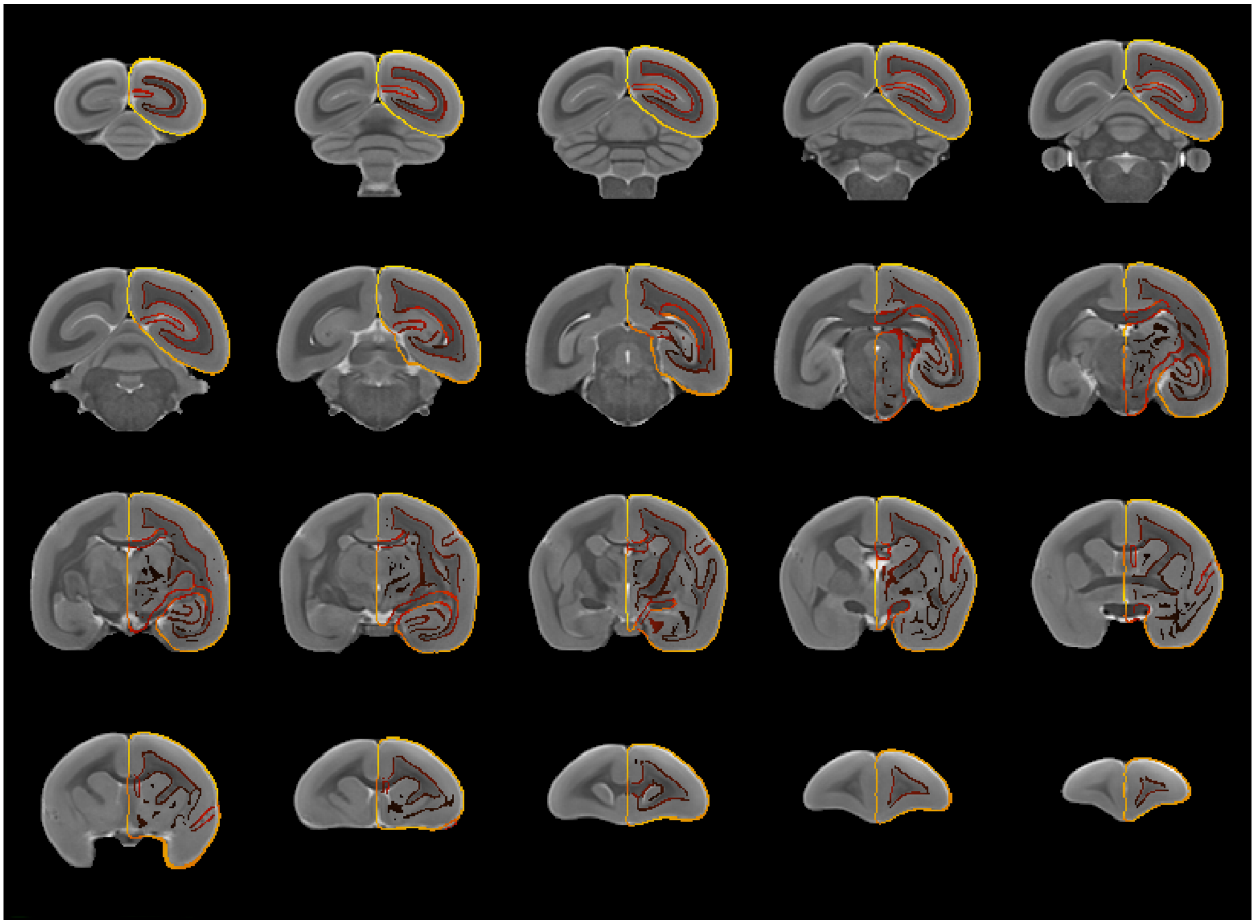

**Supplementary Figure 12. The registration of the histological NM template to the MBMv3 MRI template.** The underlay is the T2w template of the MBMv3, and the overlay is the outline of the histological NM template that is transformed on the MBMv3 template space. The outline is generated by the @AddEdge function of the AFNI (using the default setting).

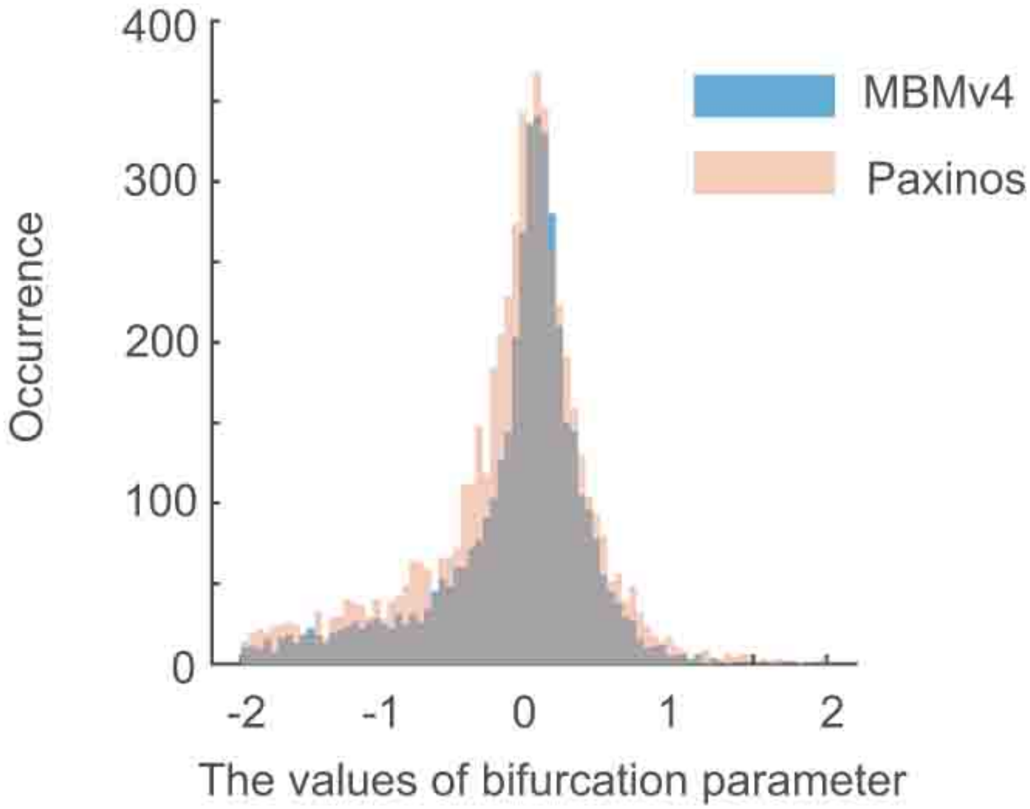


**Supplementary Figure 13. The distribution histogram of optimal bifurcation parameter.** Blue: based on MBMv4 atlas; Light red: based on Paxinos atlas;

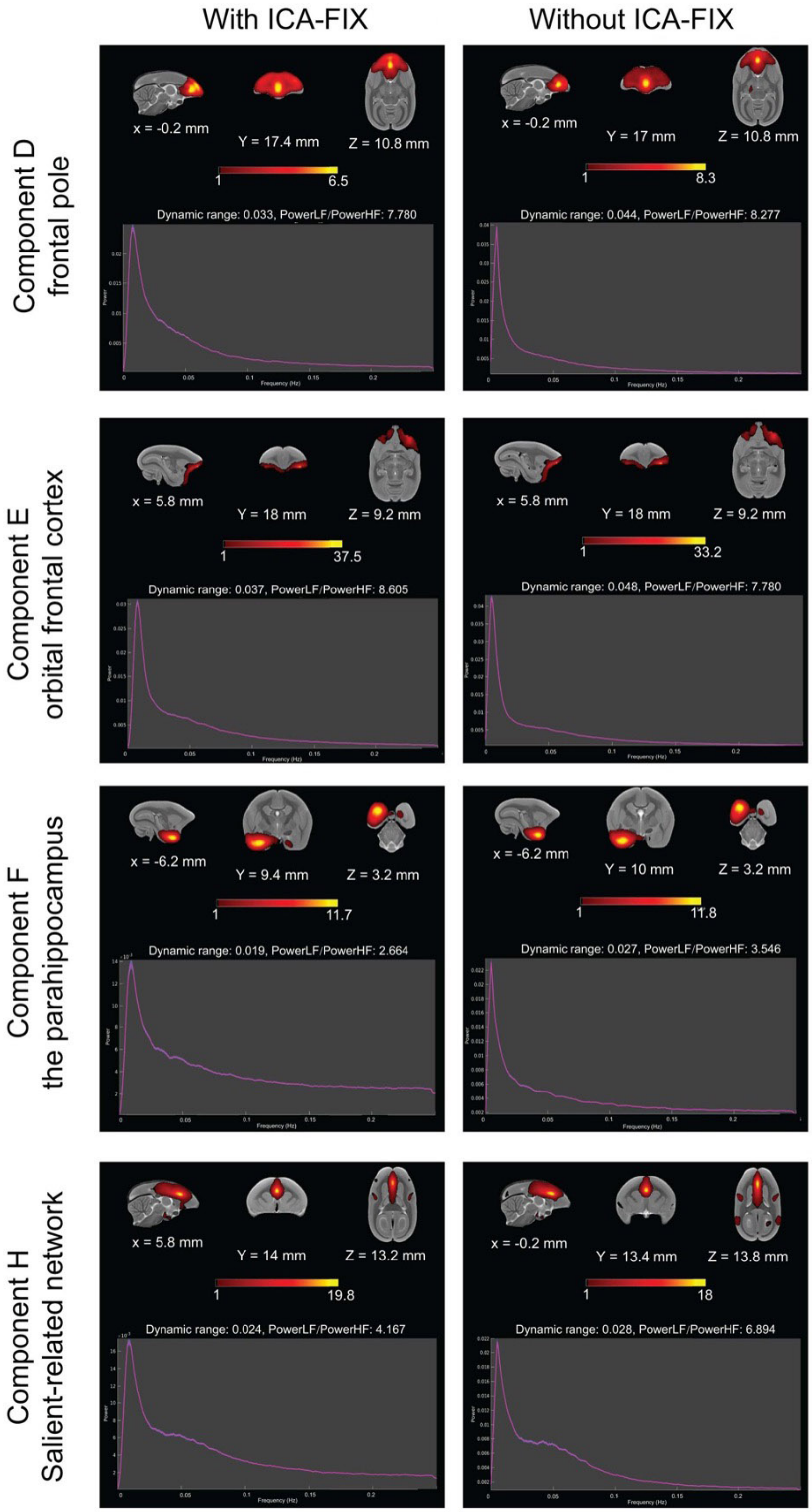


**Supplementary Figure 14. The influence of ICA-FIX on group-ICA results.** We created the ICA-FIX version of preprocessed data and ran the gICA on data with the ICA-FIX cleaning dataset. These components are similar, regardless of ICA-FIX.

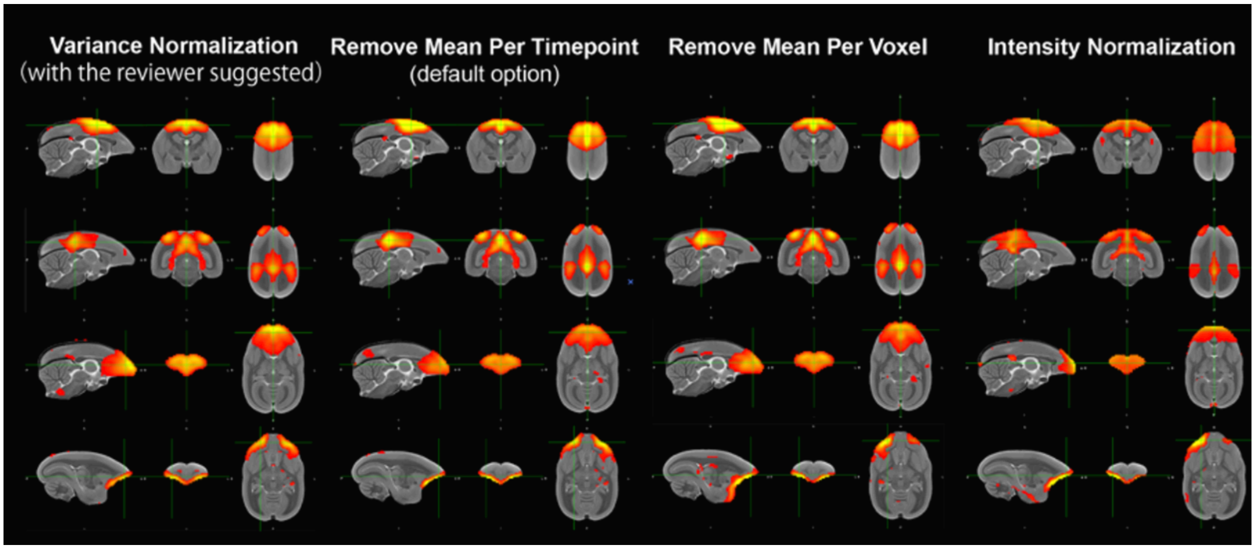


**Supplementary Figure 15. The influence of data normalization on group-ICA results.** We tested different data normalization approaches and the group-ICA results are similar.

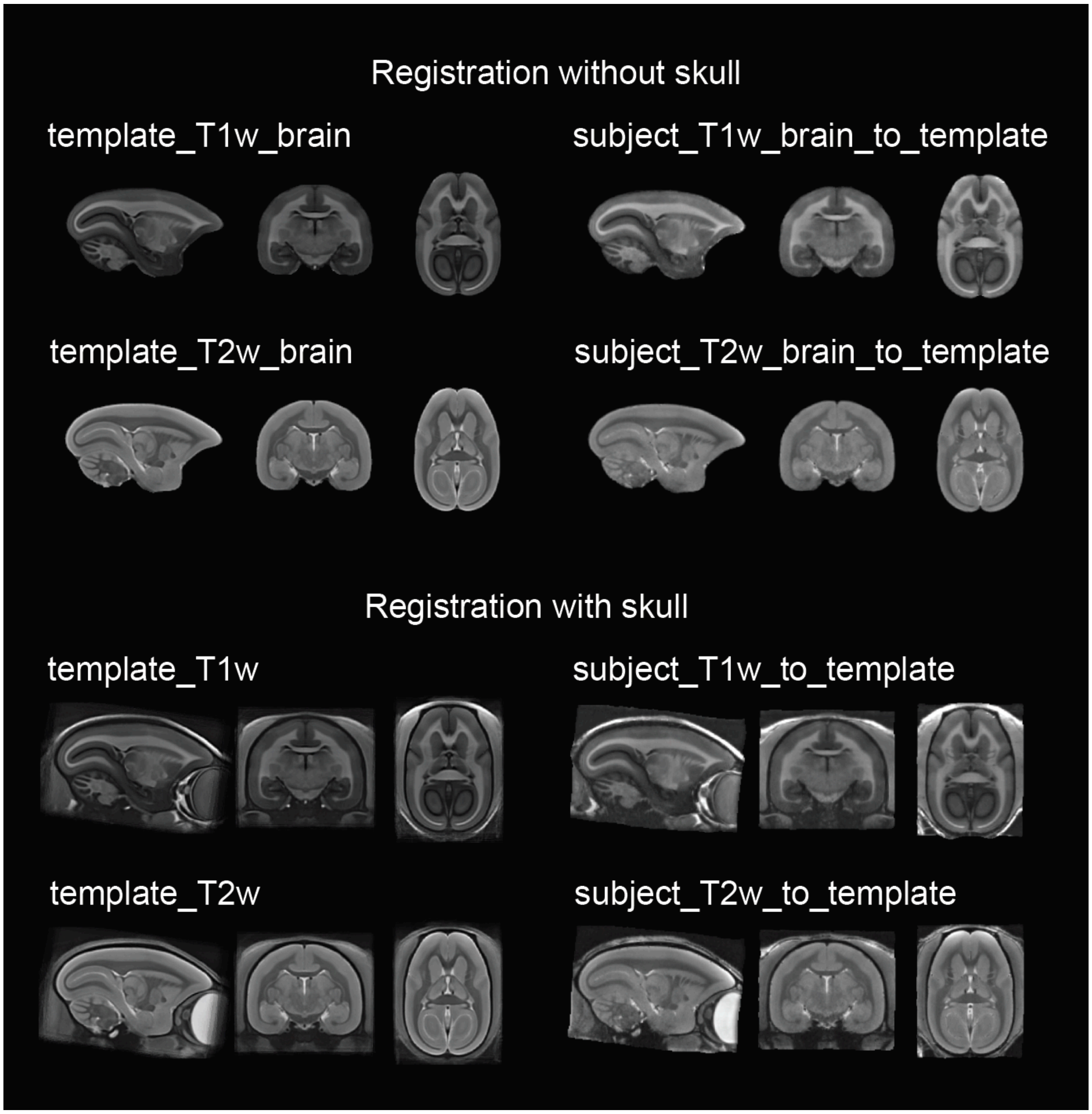


**Supplementary Figure 16. Registration results of a T1w image and a T2w image.** We tested the registration results using a T1w image and a T2w image and obtained similar results.

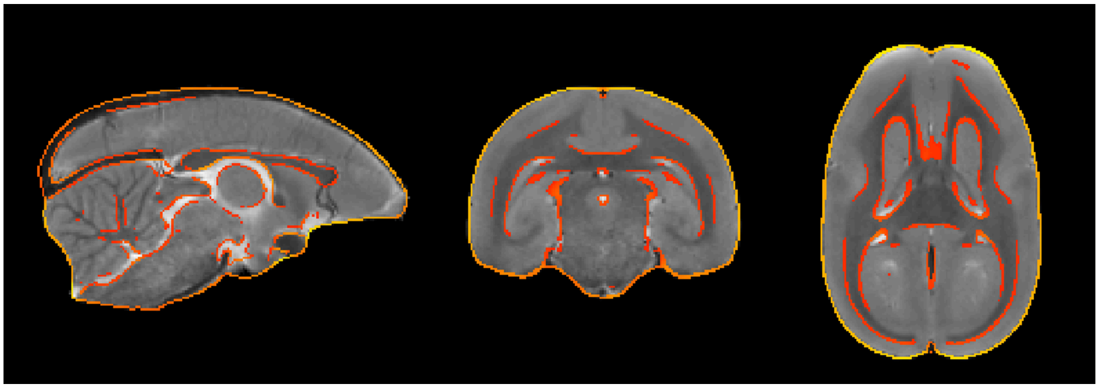

**Supplementary Figure 17. Registration results using T2w image.** The underlay is the subject T2w image transformed in the MBMv3 space, and the overlay is the outline of the T2w template of the MBMv3. The outline is generated by the @AddEdge function of the AFNI (using the default setting).

### Supplementary Tables

**Supplementary Table S1. The additional information of marmosets and datasets.** The table is provided as a Excel sheet and included in the source data, as well as the resource website (https://marmosetbrainmapping.org/data.html) under “Notes-Animal information”.

**Supplementary Table S2. Summary of head motion quality control for each fMRI run.** The table is provided as a Excel sheet and included in the source data, as well as the resource website (https://marmosetbrainmapping.org/data.html) under “rsfMRI: Head Motion Summary”.
